# Detection dogs as a help in the detection of COVID-19 Can the dog alert on COVID-19 positive persons by sniffing axillary sweat samples ? Proof-of-concept study

**DOI:** 10.1101/2020.06.03.132134

**Authors:** Dominique Grandjean, Riad Sarkis, Jean-Pierre Tourtier, Clothilde Julien-Lecocq, Aymeric Benard, Vinciane Roger, Eric Levesque, Eric Bernes-Luciani, Bruno Maestracci, Pascal Morvan, Eric Gully, David Berceau-Falancourt, Jean-Luc Pesce, Bernard Lecomte, Pierre Haufstater, Gregory Herin, Joaquin Cabrera, Quentin Muzzin, Capucine Gallet, Hélène Bacqué, Jean-Marie Broc, Leo Thomas, Anthony Lichaa, Georges Moujaes, Michele Saliba, Aurore Kuhn, Mathilde Galey, Benoit Berthail, Lucien Lapeyre, Olivier Méreau, Marie-Nicolas Matteï, Audrey Foata, Louisa Bey, Anne-Sophie Philippe, Paul Abassi, Ferri Pisani, Marlène Delarbre, Jean-Marc Orsini, Anthoni Capelli, Steevens Renault, Karim Bachir, Anthony Kovinger, Eric Comas, Aymeric Stainmesse, Erwan Etienne, Sébastien Voeltzel, Sofiane Mansouri, Marlène Berceau-Falancourt, Brice Leva, Frederic Faure, Aimé Dami, Marc Antoine Costa, Jean-Jacques Tafanelli, Jean-Benoit Luciani, Jean-Jacques Casalot, Lary Charlet, Eric Ruau, Mario Issa, Carine Grenet, Christophe Billy, Loic Desquilbet

**Affiliations:** Ecole Nationale Vétérinaire d’Alfort, Université Paris Est, 7 avenue du général de Gaulle, 94700 MAISONS-ALFORT (FRANCE); Université Franco-Libanaise St Joseph, Faculté de Médecine, Rue de Damas, BEIROUT 11072180 (LIBAN); Hôpital d’Instruction des Armées Bégin, 69 avenue de Paris, 94160 SAINT-MANDÉ (FRANCE); Service d’Incendie et de Secours de Corse du Sud (2a), Chemin de la Sposata, 20189 AJACCIO (FRANCE); Centre Hospitalo-Universitaire Henri-Mondor, Université Paris Est-Créteil, 51 avenue du Maréchal de Lattre de Tassigny, 94010 MAISONS-ALFORT (FRANCE); DiagNoses, Ferme St Grégoire, 51310 NEUVY (FRANCE); Service Départemental d’Incendie et de Secours de Seine et Marne (77), 56 avenue de Corbeil, 77001 MELUN (FRANCE); Compagnie cynophile de la Préfecture de Police, Avenue du l’Ecole de Joinville, 75012 PARIS (FRANCE); Centre Hospitalier, Route du Stiletto, 20090 AJACCIO (FRANCE); Hôpital Impératrice Eugénie, Boulevard Pascal Rossino, 20000 AJACCIO (FRANCE); Cynopro Dectection Dogs, Carrefour Porte de Madrid Pavillon est, 75016 PARIS (FRANCE); Gendarmerie Nationale Groupe d’Intervention Cynophile, Quartier Battesti CS 80402, 20162 AJACCIO (FRANCE); Grand hôpital de l’Est Francilien, 2 cours de la Gondoire, 77600 JOSSIGNY (FRANCE); Centre hospitalier François Quesnay, Boulevard de Sully, 7 avenue du général de Gaulle, 78201 MANTES-LA-JOLIE (FRANCE); Hôpital Universitaire Pitié-Salpêtrière, 47-83 Boulevard de l’Hôpital 75013 PARIS (FRANCE); Biodesive SAS, 25 rue Becquerel, 67200 STRASBOURG (FRANCE)

**Keywords:** COVID-19, SARS-CoV-2, Volatile organic compounds, Detection dogs, Olfactory detection, sweat

## Abstract

The aim of this study is to evaluate if the sweat produced by COVID-19 persons (SARS-CoV-2 PCR positive) has a different odour for trained detection dogs than the sweat produced by non COVID-19 persons. The study was conducted on 3 sites, following the same protocol procedures, and involved a total of 18 dogs. A total of 198 armpits sweat samples were obtained from different hospitals. For each involved dog, the acquisition of the specific odour of COVID-19 sweat samples required from one to four hours, with an amount of positive samples sniffing ranging from four to ten. For this proof of concept, we kept 8 dogs of the initial group (explosive detection dogs and colon cancer detection dogs), who performed a total of 368 trials, and will include the other dogs in our future studies as their adaptation to samples scenting takes more time.

The percentages of success of the dogs to find the positive sample in a line containing several other negative samples or mocks (2 to 6) were 100p100 for 4 dogs, and respectively 83p100, 84p100, 90p100 and 94p100 for the others, all significantly different from the percentage of success that would be obtained by chance alone.

We conclude that there is a very high evidence that the armpits sweat odour of COVID-19+ persons is different, and that dogs can detect a person infected by the SARS-CoV-2 virus.

---

Canine olfactive detection has proven its efficacy in numerous situations (explosives, drugs, bank notes…) including for early diagnosis of human diseases: various cancers, alert of diabetic or epileptic people in immediate alarm of crisis (1) The Nosaïs project, conducted by the UMES (Unité de Médecine de l’Elevage et du Sport) at Alfort school of veterinary medicine (France), has been set up in order to develop the scientific approach of medical detection dogs. The occurence of the COVID-19 pandemia led the Nosaïs team to start a multicentric study on the olfactive detection of SARS-CoV-2 positive people involving:

- Alfort school of veterinary, Seine et Marne Fire and Emergency department, DiagNose, Cynopro Detection Dogs and a pool of the great Paris area hospitals
- University of Corsica, South Corsica Fire and Emergency department, Corsica Gendarmerie, Ajaccio General Hospital, Ajaccio EHPAD Eugenie Hospital, Regional Health Agency of Corsica
- French-Lebanese University of Beirout, Hotel Dieu de France Hospital (Lebanon).

Fighting such a viral outbreak requires a widespread testing, one of the key measures for tackling the pandemic. In June 2020, facing a decline of COVID-19, it is possible to say that countries that have mastered their outbreak, and were able to maintain the number of infected people low, need to perform fewer test to correctly monitor the outbreak, than those countries where the virus has spread more widely. And for the same reasons, the timing of testing is also crucial. A high rate of testing will be way more effective to slow an outbreak if conducted earlier on, at a time when there is fewer infectious (2).

For this reason, and because of the potential risk of second wave of COVID-19 in the North Hemisphere, and facing the spreading of the disease in some of the South Hemisphere countries, we decided to launch the Nosaïs COVID-19 assessment on May 1^st^ 2020.

The proof-of-concept study is based on the preliminary assumption that dogs, with their highly advanced sense of smell, could be trained to discriminate COVID-19 positive people from negative ones, in relation with the excretion of specific catabolites induced by the SARS-CoV-2 virus cellular actions or replications in the sweat. Our first and main basic question, though, is “Does the sweat of a COVID-19 positive patient have a specific odour, distinct from the one of COVID-19 negative people, for trained detection dogs ?”

## Medical detection dogs: state of play

### a) Volatil Organic Compounds (VOCs)

Volatil Organic Compounds (VOCs) represent a large range of stable chemical products, volatile at an ambient temperature, and are detectable, for those of interest in dog’s olfaction, in breath, urine, tears, saliva, faeces and sweat. Most studies on volatile biomarkers have been conducted on breath samples, even if other matrices (urine and faeces mainly) have been studied.

They are more and more frequently used for dog olfactive detection of explosives, drugs, banknotes, conservation (reintroduction and survey of wild species), endangered species traffic and so on (3). Skin is the largest human organ, accounting for approximately 12 to 15 p100 of the body weight (4). VOCs emanating from skin contribute to a person’s body odour, and may convey important information about metabolic processes; they are produced by eccrine, apocrine and sebaceous glands secretions, are the major source of underarms odorants, and play a role in chemical signalling (5); on this point, numerous studies have been performed in order to characterize the specificity of underarms VOCs produced by apocrine glands (6, 7, 8, 9).

Tracking dogs trained to recognize an individual’s scent on garments from a particular body (e.g. chest, arm, underarm, leg) are not reliably able to generalize the individual’s odour to other body parts, suggesting that a different odorant milieu may characterize the back and the forearm (10).

Sweat from the palms of hands, soles of feet and the forehead comes mainly from eccrine glands and sebum; in that case, most of the compounds have been documented to be organic acids ranging in carbon size from C2 to C20, the most abundant being C16 and C18 saturated, monounsaturated and di-unsaturated, which are not volatile at body temperature (11). A more recent study confirmed that human sweat is different depending on the anatomic site on humans, and that underarm sweat appears as the most efficient in term of organic compounds turning volatile at ambient temperature close to bodylike temperature (12).

This partly explains that in our detection dogs study we chose to sample underarm sweat in the armpits.

### b) Non infectious diseases detection dogs

For centuries, our human sense of smell has been used by medicine practitioners, be it for recognising gas gangrene on the battle field or diabetic ketoacidosis. Scent detection by animals has been published in a large number of diagnostic studies, which all suggest similar or even superior accuracy compared with standard diagnostic methods (13).

A bibliography request on PubMed including the key words “dog, detection and cancer” leads to 2612 publications, demonstrating that the dog’s nose is now considered as a tool for early detection of main cancers and their prevention. An exhaustive review was published by Angle in 2016 (14) on the subject.

The hypothesis that dogs could be able to detect malignant tumors based on their specific odour was put forward by Williams in 1989 (15). The first clinical investigation of cancer was published by Willis

(16) on bladder cancer in 2010, after having published a proof of principle study in 2004 (17). Several positive studies have been performed on colorectal cancer screening with odour material by canine scent detection (18, 19), and some tendencies have been recorded for lung cancer (20, 21, 22, 23), melanoma (24, 25), prostate (26, 27, 28), or liver (29) cancers.

The dog’s nose is also now currently utilised in order to prevent the consequences of crisis (or even prevent them) for diabetic (30, 31, 32, 33) and epileptic people (70).

New diseases are starting to be explored (ovarian cancers or degenerative diseases like Parkinson), as dogs training techniques are now much better known and specific volatile organic compounds considered as solid biomarkers.

### c) Infectious and parasitic diseases

Several studies have used dogs in attempt to differentiate a range of target odours among insects and parasites. For example, Wallner (34) trained dogs to locate egg masters of the gypsy moth with a 95p100 success rate. A study conducted by Richards (35) assessed the dog ability to differentiate between nematods infected and uninfected sheep faeces. But recently, Guest (36) did show that trained dogs can identify people infested with malaria parasites thanks to their olfactive capacities.

Bacteriological diseases can also be detected by the dog’s superior olfactory system as Bomers (37, 38) demonstrated during a hospital outbreak of Clostridium difficile. Rats’ olfaction is also used now for the detection of tuberculosis in East Africa (39).

In a study by Maurer (40) dogs were trained to distinguish urine samples that were positive for bacteriuria (*E. Coli, Enterococcus, Staphylococcus aureus, Klebsiella*) from those of culture-negative controls. All dogs performed with similarly accuracy: overall sensitivity was at or near 100p100, and specificity was above 90p100.

Concerning virus detection, dogs are able to differentiate cellular cultures infected by the virus of the bovine mucosae disease from non-infected cultures, or from those infected by other viruses (bovine herpes 1 or bovine parainfluenza 3) (14).

The knowledge we have now, with a permanent increase in scientific backgrounds, led the Nosaïs canine medical detection team of Alfort School of veterinary medicine to set up a program related to the early detection of COVID-19 on humans. The first phase of the program is to train dogs to sniff human sweat samples and see if they can differentiate the sweat odour of COVID-19+ patients compared to that of COVID-19-persons. We make the assumption that the replications and cellular actions of the virus SARS-CoV-2 generate specific metabolites and catabolites that can be excreted by the apocrine sudoral glands and generate VOCs that the dogs can “identify”. In a second time, we attempt to validate the method with the dogs checking samples one by one instead of comparatively, in order to fit with what could be an operational utilisation of such trained dogs.

## Material and Methods

The study is conducted on 3 different sites following the same protocol:

- Paris (France): Alfort School of Veterinary Medicine
- Ajaccio (France): South Corsica Fire and Emergency Dept
- Beirout (Lebanon): French-Lebanese University Saint Joseph

### a) Samples

#### Nature of samples

We chose to work on axillary sweat samples for different reasons:

- Those already explained in a previous paragraph of this paper
- Sweat does not appear to be a route of excretion for SARS-CoV-2 virus (41, 42)
- Anticipating possible future practical applications of dogs trained to detect people carrying the SARS-CoV-2 virus; if the results of our studies are positive, sweat (even if not the only one) is the key odour for search and rescue or tracking dogs (43)
- Anatomical sites of sampling are armpits, considering this site is hardly passively contaminable by the positive patient (Figure 1)

**Figure 1:**
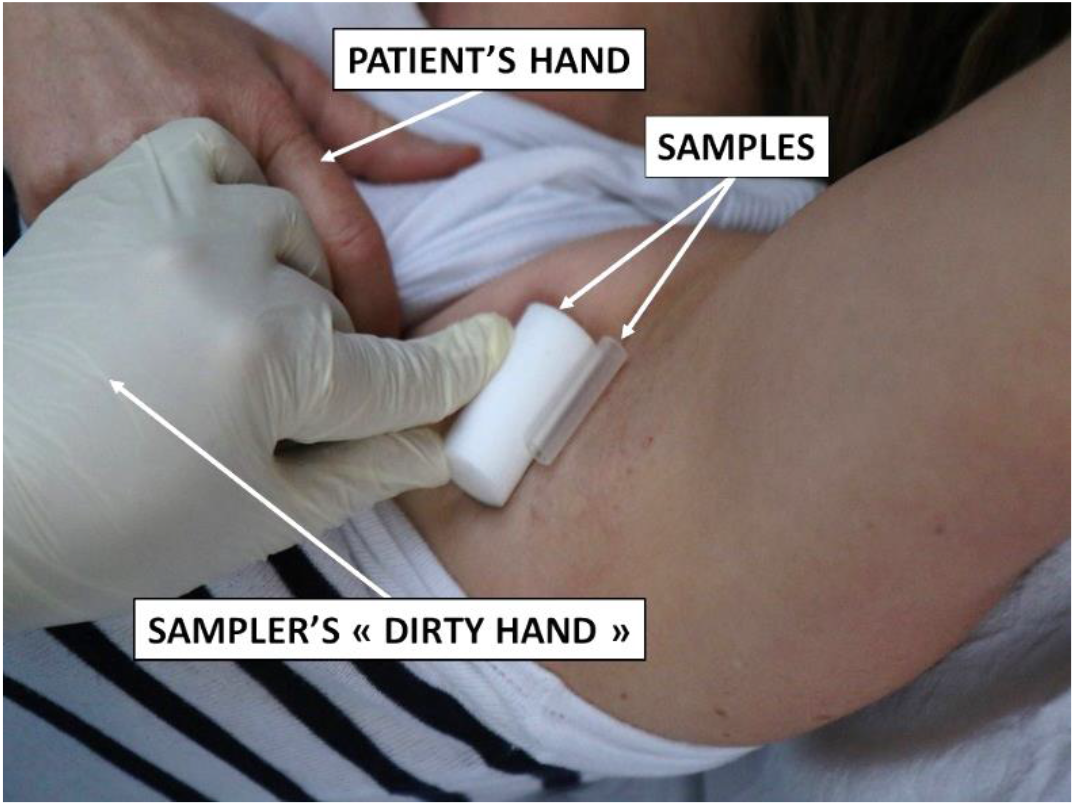
Sampling in armpit

#### Inclusion and exclusion criteria

Positive samples are realised on patients showing clinical symptoms of COVID-19, who were COVID-19 positive on RT-PCR or PCR test for SARS-CoV-2.

Patients were excluded from the study protocole if they had received a medical treatment for more than 36 hours, in order to avoid possible interferences with long-term medical treatments in the sweat VOCs. Samples are collected by doctors, interns or nurses who were trained to not contaminate the samples with their own odours.

A training video and the necessary equipment has been provided to each staff in charge of samples collections.

Negative samples are collected the same wa, by the same trained staffs, from patients who met the following inclusion criteria:

- no clinical symptoms related to COVID-19
- negative on SARS-CoV-2 PCR test

In order to avoid any dog creancement on the olfactive “background noise” of positive sampling sites, both positive and negative samples collections were made in the same locations.

A total of 198 samples (101 positive samples and 97 negative samples) were collected at the endpoint date of the present study (May 28^th^, 2020), and their distribution according to sites and sex is reported on Table 1.

**Table 1:** Distribution of the axillary sweat samples realised on each site

| Samples | Paris |  |  |  | Ajaccio |  |  |  | Beirout |  |  |  |
| --- | --- | --- | --- | --- | --- | --- | --- | --- | --- | --- | --- | --- |
|  | + |  | - |  | + |  | - |  | + |  | - |  |
|  | ♀ | ♂ | ♀ | ♂ | ♀ | ♂ | ♀ | ♂ | ♀ | ♂ | ♀ | ♂ |
| 0-19 years old | 0 | 0 | 1 | 2 | 0 | 0 | 0 | 2 | 1 | 2 | 0 | 1 |
| 20-39 years old | 1 | 0 | 11 | 4 | 0 | 0 | 2 | 5 | 15 | 16 | 8 | 7 |
| 40-59 years old | 4 | 3 | 4 | 4 | 0 | 0 | 1 | 5 | 12 | 14 | 11 | 10 |
| 60-79 yeas old | 2 | 7 | 2 | 2 | 1 | 0 | 0 | 0 | 1 | 7 | 2 | 8 |
| ≥ 80 years old | 7 | 2 | 1 | 1 | 5 | 0 | 0 | 0 | 0 | 0 | 1 | 0 |
| Unspecified | 1 |  | 1 |  |  |  |  |  |  |  |  |  |
| Total | 27 |  | 34 |  | 6 |  | 15 |  | 68 |  | 48 |  |

#### Sampling supports

Two types of sampling supports are used:

- Sterile gauze swabs utilised by the hospitals or sterile gauze filters utilised for explosive detection (provided by DiagNose comp.)
- Inert polymers tubes utilised for explosives, drugs or criminology detection (provided by Gextent comp.)(Figure 2)

**Figure 2:**
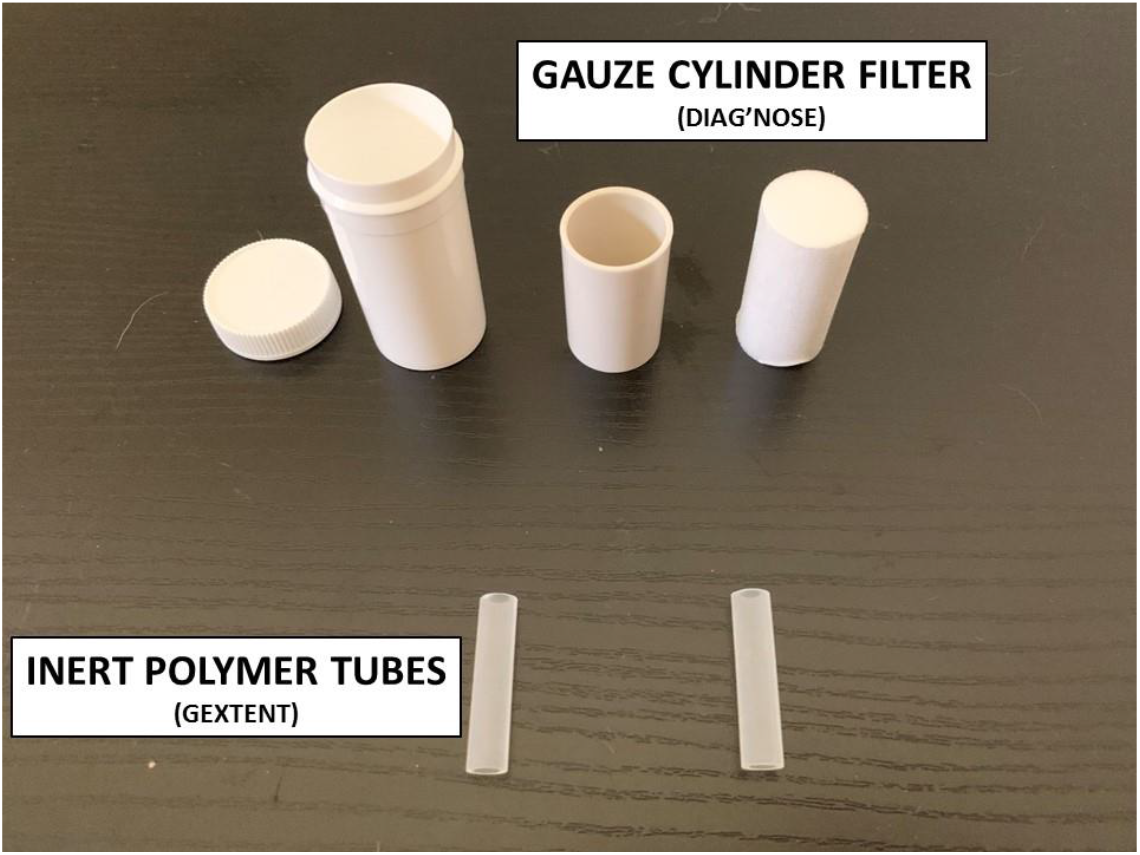
Sampling supports

The polymers tubes are able to adsorb both polar and apolar molecules, and therefore will be sent to the University of Corsica Pascal Paoli (France) for precise chemical analysis in order to try to identify sweat biomarkers of COVID-19.

#### Practical realisation and storage of samples

All samples are taken in different hospitals (Great Paris, Ajaccio, Beirout) by medical doctors or internists, helped by a nurse.

Negative samples do not request particular safety protection, the sampler wearing a new pair of surgical gloves in order not to contaminate the sample with his own odours.

Positive samples are realised wearing the full COVID-19 safety protection and two pairs of new gloves for the sampler.

Samples are put in anti-UV sterile containers, disinfected by the helper, coded (including Left or Right armpit), and those then placed in a second plastic envelope.

A sample is never used for dog training prior 24 hours after sampling the sweat, as the SARS-CoV-2 virus does not survive more than a few hours on cotton and disposable gauze (44). A more recent study concludes that absorbent materials like cotton and gauze are way safer than no-absorptive materials for protection from SARS-CoV-2 infection (45).

All coded samples are then stored at a constant temperature (18°C) and hygrometry (45p100).

Before being sampled, each person receives explanations on the protocol and signs an individual enlightened agreement form that stays by the hospital in the medical files of the patients. (Figure 3)

**Figure 3:**
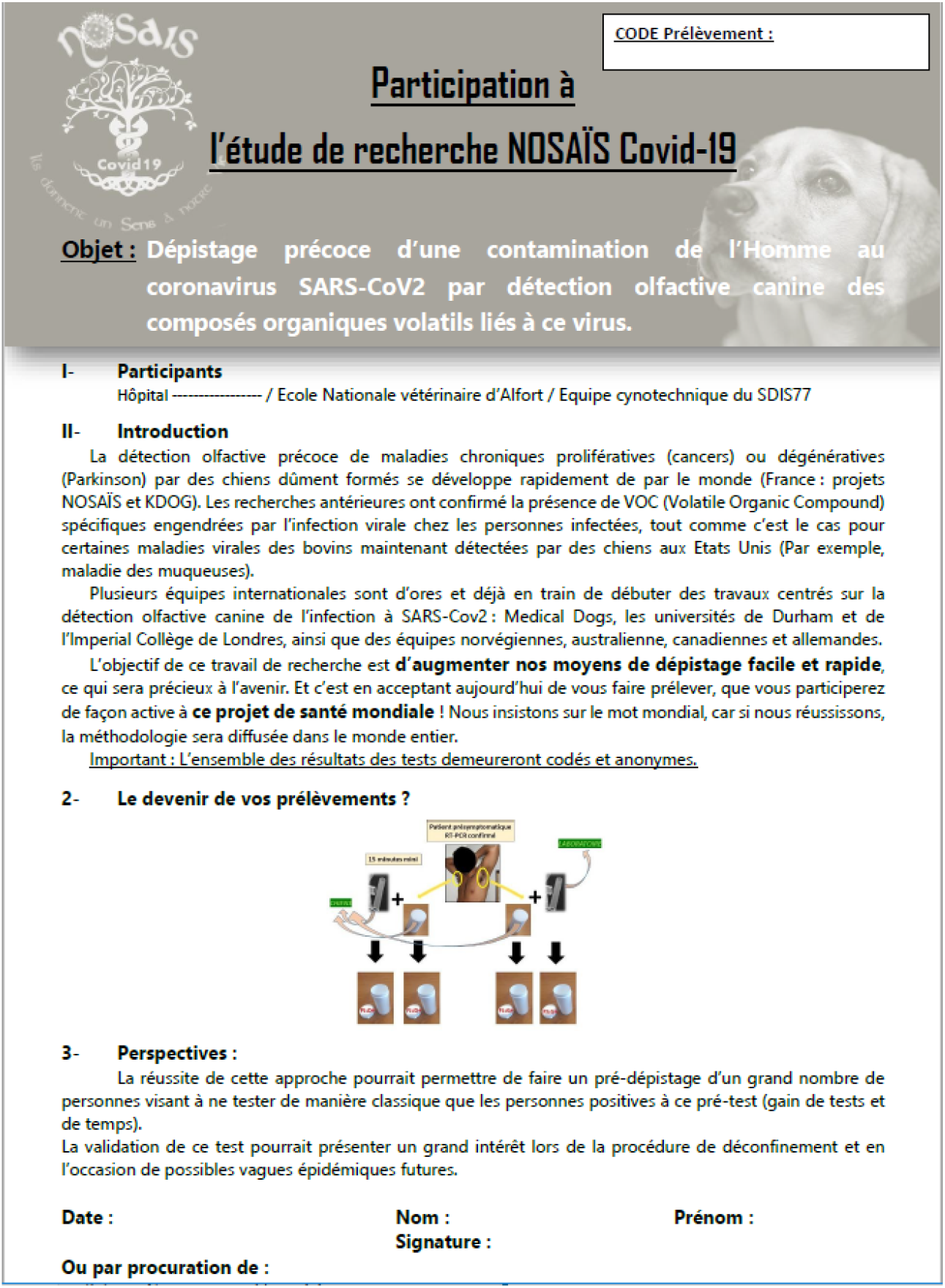
Individual enlightened agreement form (in french as provided to people)

Individual anonymous data are registered on a second form that (table 2) follows the coded samples.

**Table 2:**
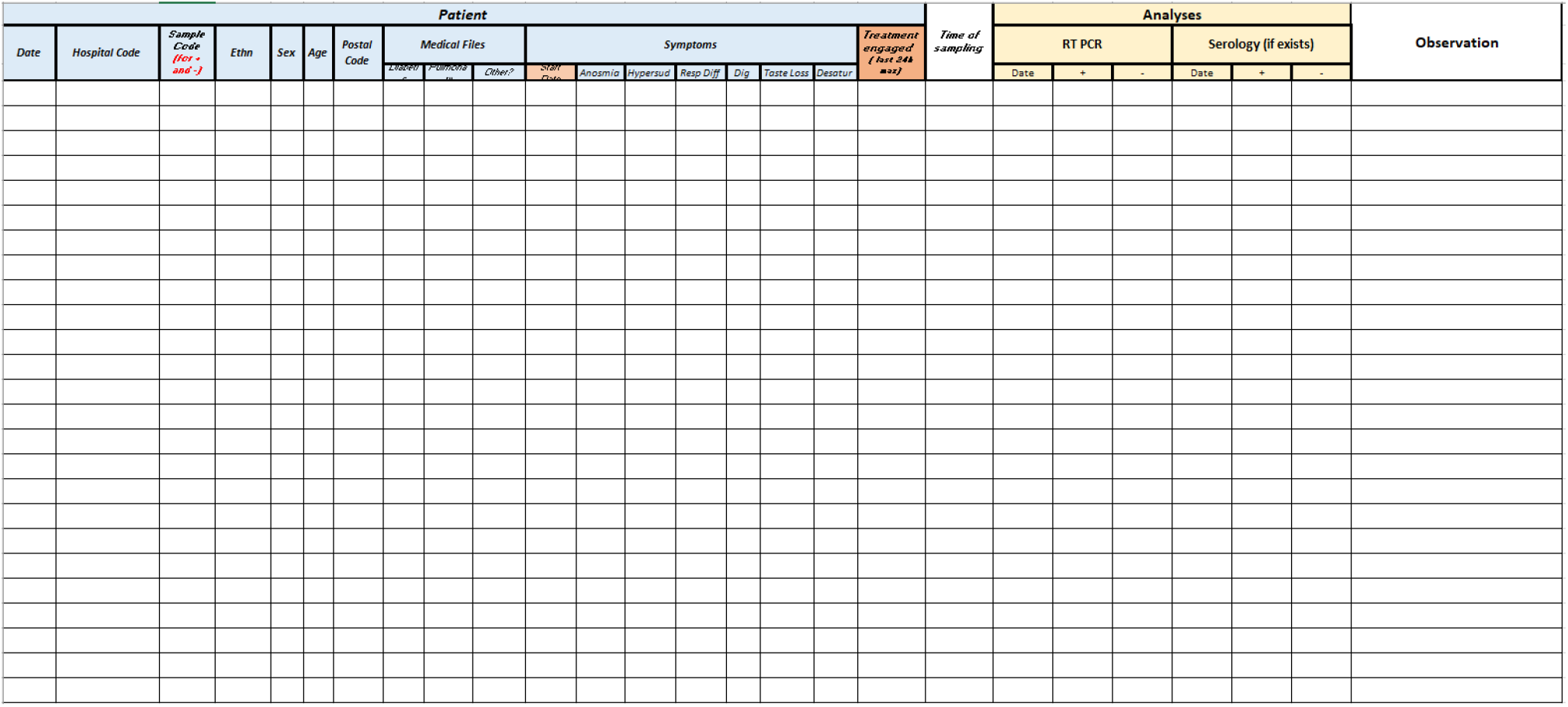
Anonymous sampled persons data registration form?

In Beirout (Lebanon), due to the local hot climate, samples are stored at a temperature of +6°C, constant hygrometry.

### b) Canine Ressources

#### The dogs

The 18 dogs that started the experimentation and their characteristics are regrouped in Table 3 and belong to:

**Table 3:** Characteristics of the dogs entering the study

|  | Name | Gender | Breed | Age | Structure involved | Specialty |
| --- | --- | --- | --- | --- | --- | --- |
| Paris | Guess | F | Belgian Malinois | 8 years | Cynopro detection Dog | Explosives |
|  | Gun | M | Belgian Malinois | 16 months | Cynopro detection Dog | Explosives |
|  | Bruce | M | German shepherd | 2 years | DiagNose | Explosives |
|  | Oslo | M | Belgian Malinois | 18 months | DiagNose | Explosives |
|  | Diablo | M | Belgian Malinois | 12 years | Fire and Emerg. Dept. 77 | Search and rescue |
|  | Fidjie | F | Belgian Malinois | 6 years | Fire and Emerg. Dept. 77 | Search and rescue |
|  | Jet | M | Belgian Malinois | 6 years | Fire and Emerg. Dept. 77 | Search and rescue |
|  | Joy | F | Belgian Malinois | 6 years | Fire and Emerg. Dept. 60 | Arson |
|  | Lesko | M | Belgian Malinois | 5 years | Fire and Emerg. Dept. 77 | Search and rescue |
|  | Loki | M | Belgian Malinois | 5 years | Fire and Emerg. Dept. 77 | Search and rescue |
|  | Maïka | F | Belgian Malinois | 3 years | Fire and Emerg. Dept. 77 | Search and rescue |
|  | Phoenix | M | Belgian Malinois | 15 months | Fire and Emerg. Dept. 77 | Search and rescue |
| Ajaccio | Asko | M | Belgian Malinois | 3 years | Fire and Emerg. Dept. 2a | Search and rescue |
|  | HK | F | Belgian Malinois | 8 years | K9 Invest. Group Gendarmerie of Corsica | Drugs, Banknotes |
|  | M'Inty | M | Belgian Malinois | 3 years | Fire and Emerg. Dept. 2a | Search and rescue |
|  | Nikita | F | Belgian Malinois | 2 years | Fire and Emerg. Dept. 2a | Search and rescue |
| Beirout | Bella | F | Belgian Malinois | 6 years | Mario-K9 | Colon Cancer |
|  | Jacky | M | Jack Russell terrier | 3 years | Mario-K9 | Colon Cancer |

- Paris site: Firefighters and Emergency Dept. 77 (Seine et Marne), DiagNose comp., CynoPro Detection Dog comp.
- Ajaccio site: Firefighters and Emergency Dept. 2a (South Corsica), National Gendarmerie (Corsica)
- Beirout site: Barbara (Lebanon) Mario k9

We just present two of the dogs who participates to our study (Figures 4 and 5)

**Figure 4:**
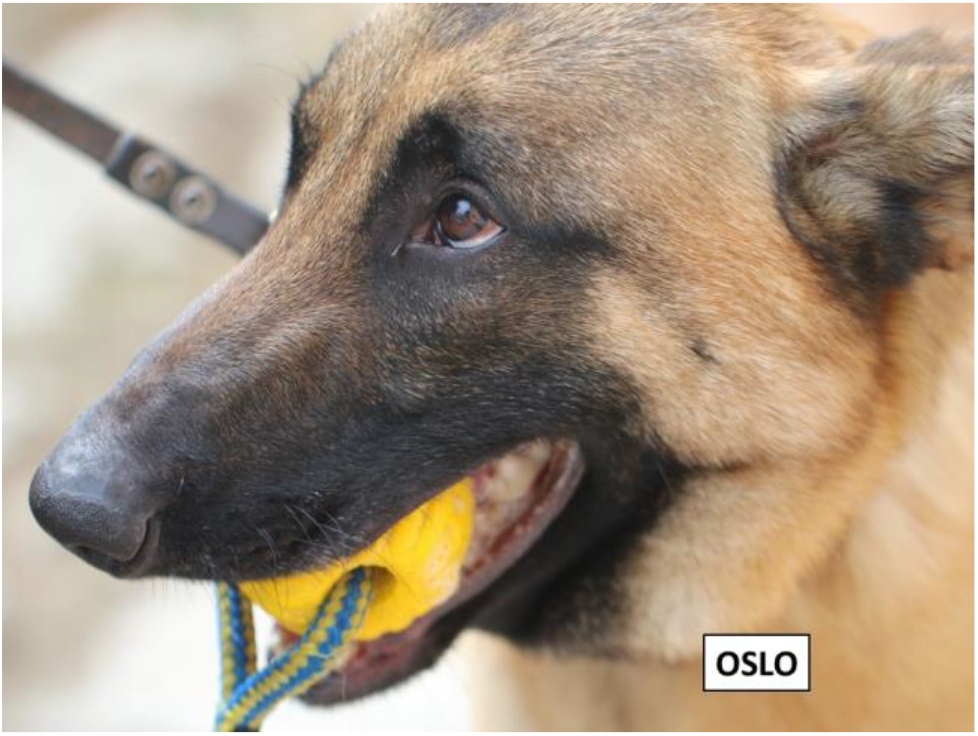
Dog “Oslo”

**Figure 5:**
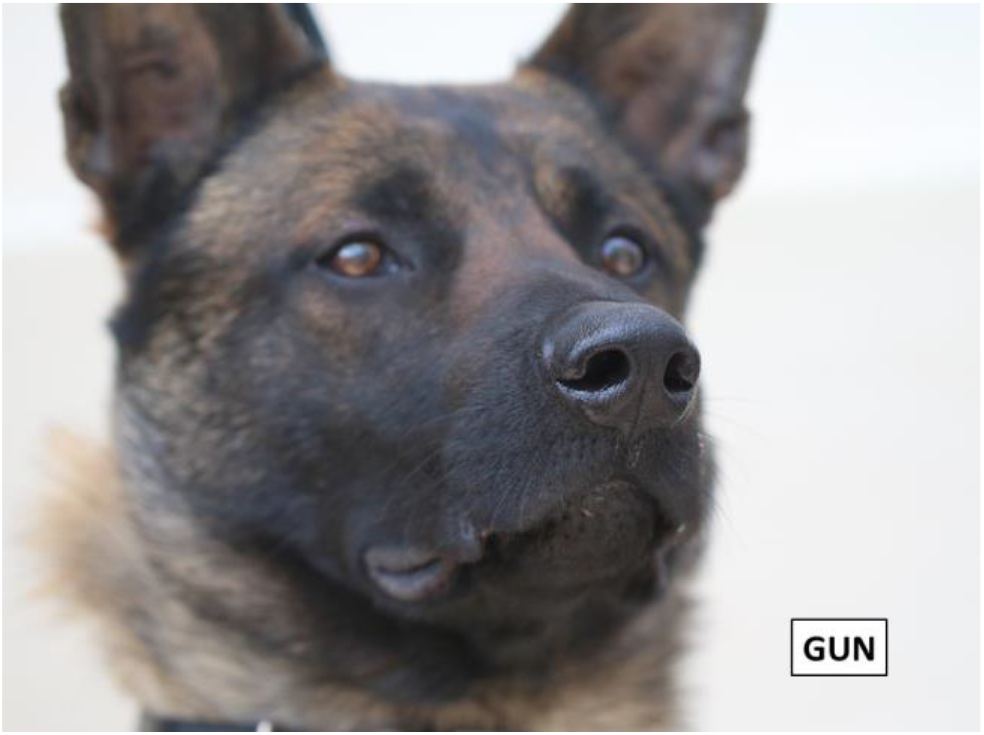
Dog “Gun”

For this study we decided to include three different types of detection dogs (Table 3):

- Explosives detection dogs, trained to detect 30 to 40 different types of explosives but moreover used to work on a line of samples that they have to sniff individually. For such dogs, if COVID-19+ samples have a specific odour, they only have to upload one more odour in their olfactive memory bank.
- Search and rescue dogs, trained to perform disaster and area search, mainly work through the scent of the sweat of the persons, which explains also our choice.
- Colon cancer detection dogs trained on rectal gases.

We did not decide to work with drug detection dogs as there is always a possibility that COVID-19 positive or negative people use prohibited substances that would let catabolites be excreted by the axillary sweat.

Most of the dogs included in our study are Belgian Malinois Shepherds because it is actually the most represented breed in working dogs french structure, but we insist on the point that lots of canine breeds or mongrel dogs could develop the same olfactive detection qualities.

#### Safety of the dogs regarding SARS-CoV-2 virus infection

There has been very few, isolated and criticizable reports on the passive carriage of SARS-CoV-2 virus by the dog, with very small amounts of viral RNA, indicating that the samples, collected by an infected person, had a very low viral load (46). The low viral titres observed in this dog suggest it had developed a low-productive infection, and the likelihood of infectious transmission was minimal or none existent.

A second case was alerted in Hong-Kong when the owner tested positive for COVID-19 infection stayed in quarantine with his two dogs. One of them was tested positive for quantitative PCR but never had any symptom, the other stayed negative (47). As with the first dog, the infection was very low positive and non-contagious.

In the USA, Idexx Laboratories tested more than 4000 canine specimens during its validation of a new veterinary test system for the SARS-CoV-2 virus and found no positive animal (48). More recently a first study conducted in Alfort School of Veterinary Medicine (France) showed the absence of SARS-CoV-2 infection in dogs in close contact of a cluster of COVID-19 patients (49).

Finally, the CDC (Center for Disease Control and Prevention) in the USA (50), and the ANSES (Agence Nationale de Sécurité Environnementale et Sanitaire) in France (51) attest that there is absolutely no evidence that pet animals, and especially dogs, play any significant role in the transmission or in spreading the virus that causes COVID-19, and the risk is considered as close to zero.

#### Training and first testing of the dogs

The dogs are first trained to work on a line of sample carriers (Figures 6 and 7), and dogs are taught to sit in front of the cone (box) containing the positive sample. The method is based on positive reinforcement (the dog gets his toy for each correct marking).

**Figure 6:**
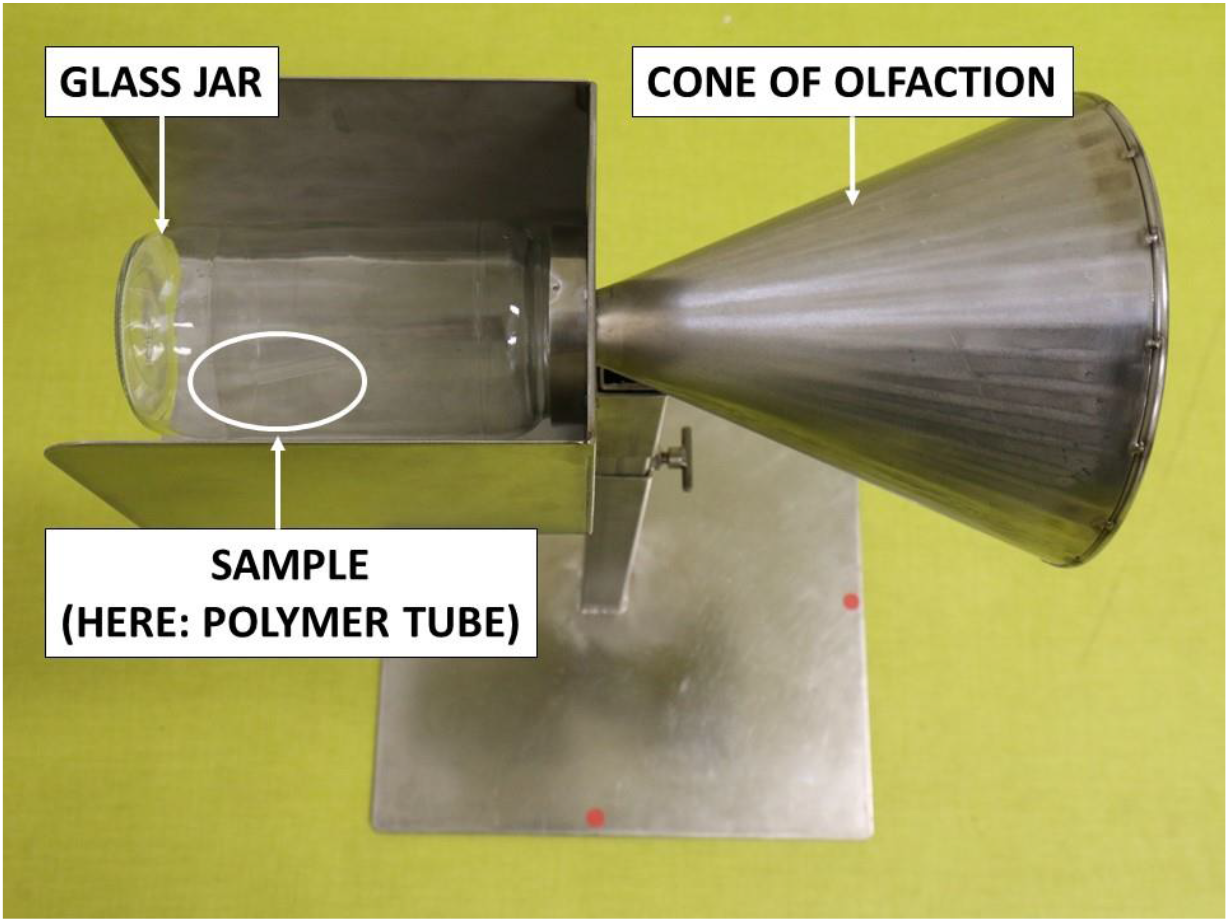
Testing equipment

**Figure 7:**
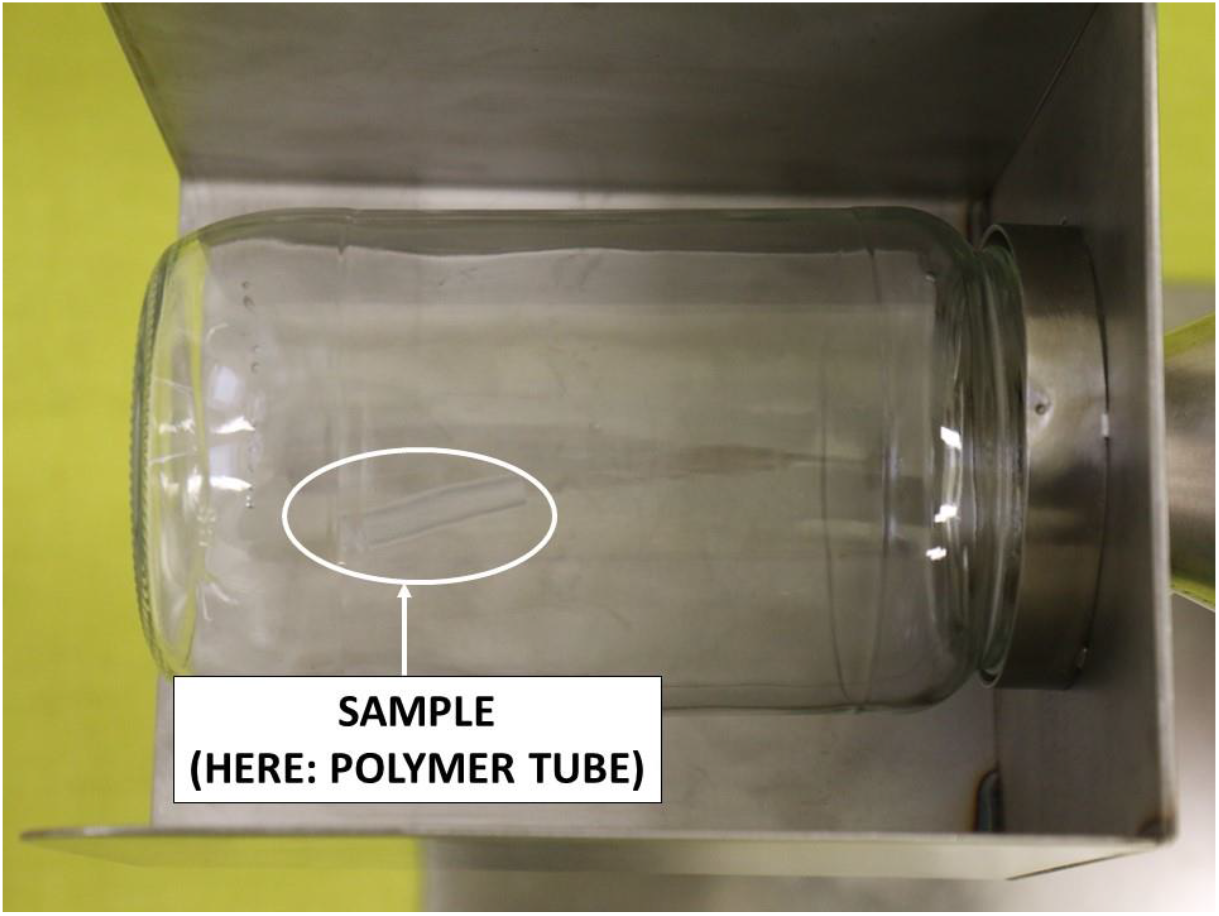
Testing equipment details

For each involved dog, the acquisition of the specific odour of COVIS-19+ sweat samples required from one to four hours, with an amount of positive samples sniffing ranging from four to ten.

#### Testing protocol

Once the training period was considered as completed by the dog’s handler, the dog was not anymore in contact with any sample prior to a trials session, and the data collection regarding the success or failure to detect a COVID19+ sweat odour started.

At that time, the dog must alert for COVID19+ sweat odour contained in one box only, out of 3,4, 6 or 7 boxes(depending on the study site, see Figure 8) presented in a line. The other boxes contained negative samples or mocks, but always with only mock in the line (Figure 9).

**Figure 8:**
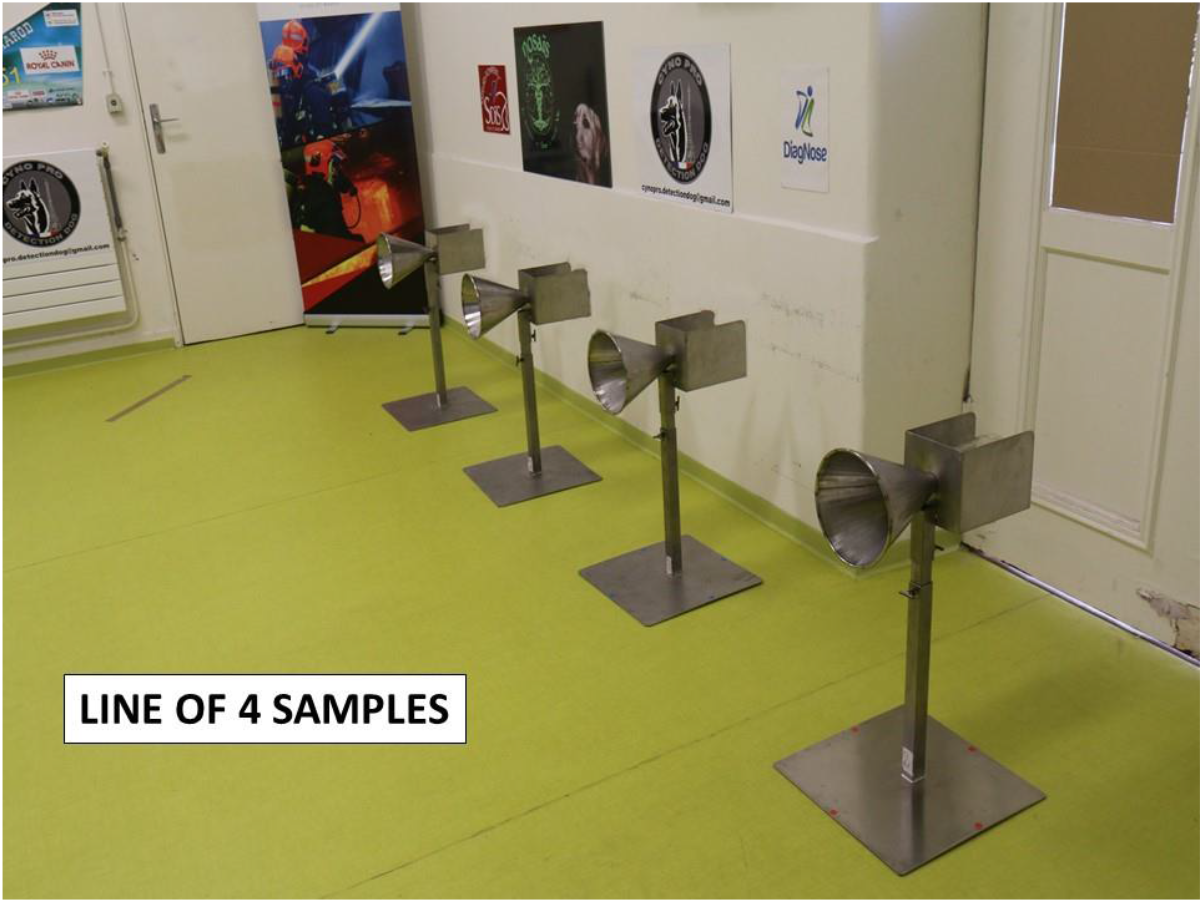
Testing line of 4 samples

**Figure 9:**
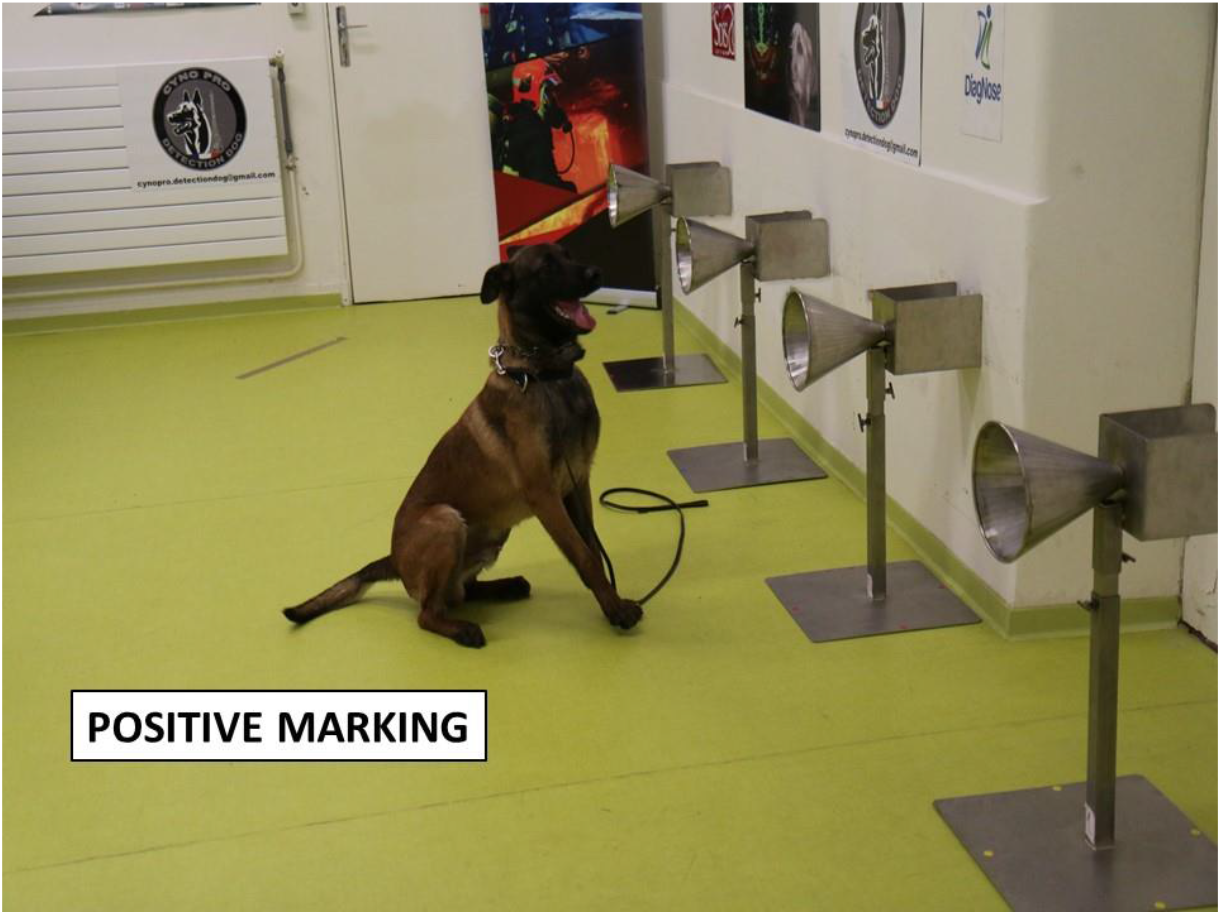
Positive marking on the line

This whole period lasted 21 days because most dogs could not work on a daily basis. No disease, symptoms or clinical abnormalities have been reported on any of the dogs involved in the study during the whole testing period.

The standard environmental conditions for canine olfactive detection were observed (temperature +18°C, hygrometry 50p100).

A “trial” consisted of one dog detecting the presence of the COVID-19+ in one box out of the 3, 4, 6 or 7 boxes. One positive sample could be used several times at different spots in the line for the same dog (3 times maximum). All the trials are independent from each other, which means that for each trial, the location of the COVID-19 positive box was randomly assigned by using a randomization dedicated wellsite (52)

Both samples and sample carriers were manipulated by the same person, wearing sanitary barrier protections and a pair of new surgical gloves at each trial, in order to avoid any olfactive contamination or interaction. All sample carriers were disinfected (acetone) between each trial.

Both the dog and his handler stayed in another room with no visual or audio access to the testing room between two trials, when the COVID-19+ sample was placed in a new box randomly assigned.

The same two researchers were dedicated to the registration of individual data (Table 4). A success was defined if the dog alerts the one box containing the COVID-19+ sweat sample (Figure 9). A failure was defined if the dog alerts on any of the other boxes. Trials were registered from the moment dogs recognized positive sample among other samples (mock or negative ones) without having sniffed any of them previously to the test. If the dog marks first a negative sample, the trial is considered as failed even if he marks the positive afterwards.

**Table 4:**
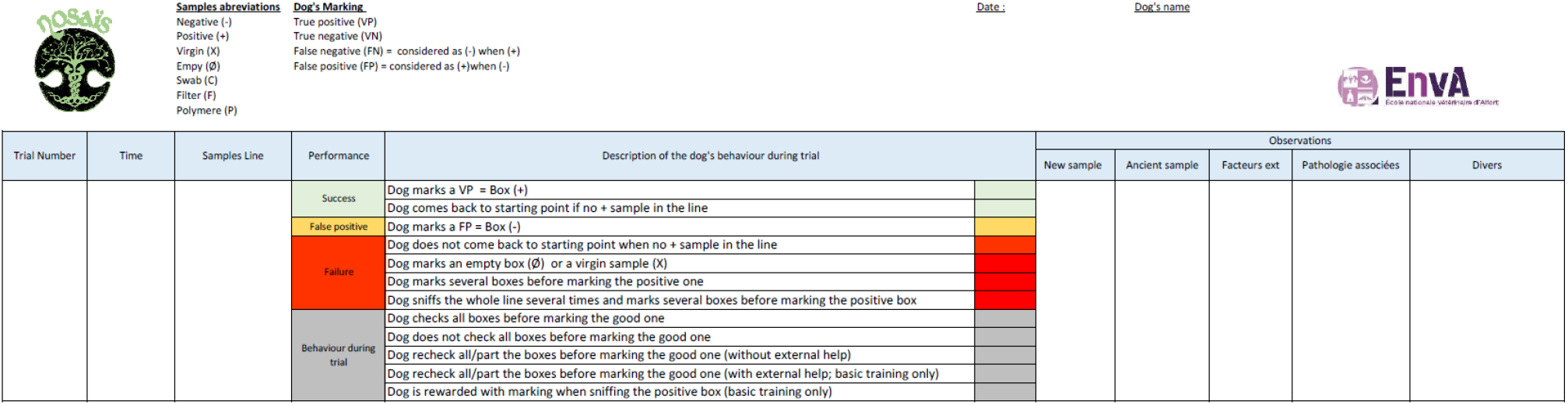
Trials individual registration form

## Results and statistical analysis

Since the objective of the study is a proof-of-concept one, we did not aim to provide evidence that a whole population of trained dogs are able to detect COVID-19 positive samples, but rather to provide evidence that a well-trained dog is able to detect COVID-19 positive samples.

Such evidence is provided if one observes from one dog a higher proportion of successes than the one expected by chance alone. Therefore, the target population is not a population of dogs, but the whole population of trials (consisting of infinitive number of trials) for one dog, each dog being considered as a “diagnostic tool”.

At the end point date, three search and rescue dogs have been withdrawn of the protocol for being unable to adapt to an olfactive search on a line of samples and eight of the same category are late in their testing period due to this necessary basic “retraining”.

The results are therefore presented on eight dogs who followed the whole process, as they appear demonstrative enough to answer our first question. A total of 232 trials were made using 33 positive samples in France (Alfort and Ajaccio), and 136 trials for 68 positive samples in Lebanon (Beirout).

In order to provide evidence that the successes were not obtained by chance alone, the proportion of successes for one dog out of the total number of trials was calculated, along with it 95p100 confidence intervals (95%CI) calculated by using the Clopper-Pearson’s method (53). The proportion of successes obtained by chance alone (thereafter called “random choice proportion”) was calculated by dividing one by the total number of sample carriers placed in the sniffing line. All the trials for one dog used the same total number of sample carriers. This total number of sample carriers depended on the site location. If the value of the random choice was not included within the 95% of the estimated proportion of successes for one dog, the null hypothesis according to which the dog chose the COVID-19 positive sample carrier by chance was rejected with a probability of type-i error of 0,05 (Table 5).

**Table 5:**
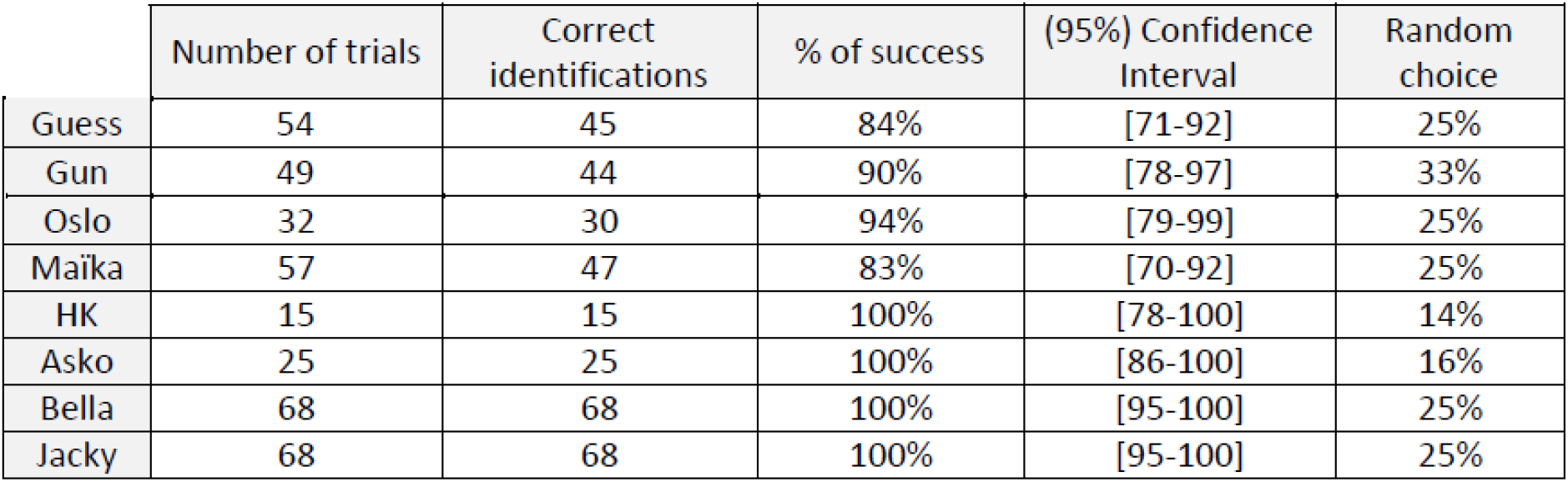
Results obtained on the dogs that finished the training period (n=8)

|  | Number of trials | Correct identifications | % of success | (95%) Confidence Interval | Random choice |
| --- | --- | --- | --- | --- | --- |
| Guess | 54 | 45 | 84% | [71-92] | 25% |
| Gun | 49 | 44 | 90% | [78-97] | 33% |
| Oslo | 32 | 30 | 94% | [79-99] | 25% |
| Maïka | 57 | 47 | 83% | [70-92] | 25% |
| HK | 15 | 15 | 100% | [78-100] | 14% |
| Asko | 25 | 25 | 100% | [86-100] | 16% |
| Bella | 68 | 68 | 100% | [95-100] | 25% |
| Jacky | 68 | 68 | 100% | [95-100] | 25% |

Table 5 presents the results obtained from the seven dogs of the study. The observed proportion of successes (i.e., the dog correctly detected the COVID-19 positive sample carrier) for each dog was significantly different from the one which would be observed by chance alone.

## Discussion

The results of this first proof of concept study demonstrate that COVID-19 positive people produce an axillary sweat that has a different odour, for the detection dog, than COVID-19 negative persons. It strongly suggests the hypothesis according to which dogs can be trained to detect SARS-CoV-2 contaminating people, contained in axillary sweat which is easy and safe to collect (Figure 10).

**Figure 10:**
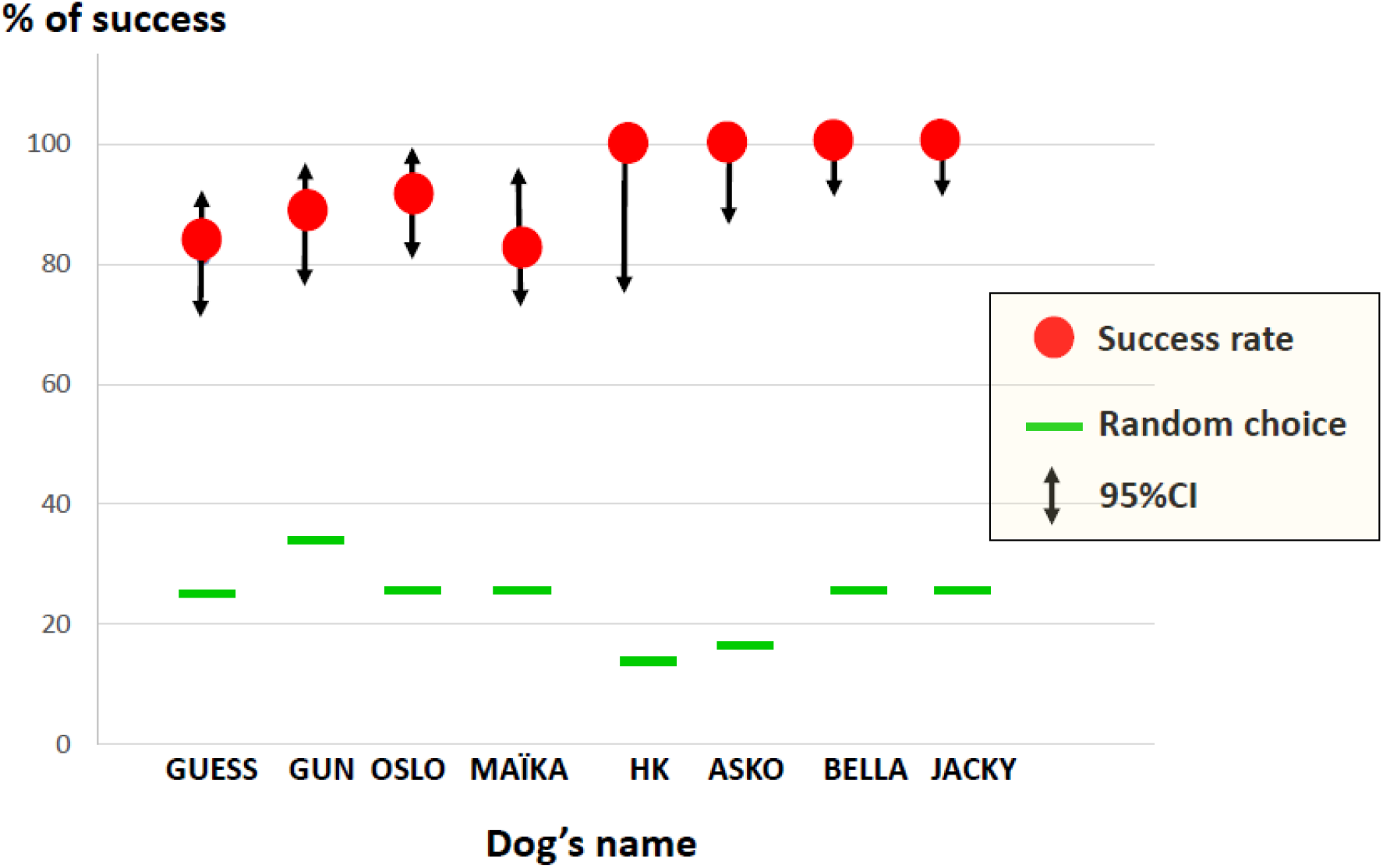
Individual results of the dogs and confidence intervals (CI) of the study and random choices values

Regardless of the nature of the dog’s detection system and the fact that biology will never be an exact science, both false positive (where the dog detects the target present when it is absent) and false negative (where the target is present but the dog fails to detect it) errors occur in every single detection system, including highly sophisticated machines.

Uncertain negative results of RT-PCR regarding COVID-19 range from 22 to 30p100 (54, 55, 56), and RT-PCR results from several test at different points appear variable from the same patients during the course for diagnosis and treatment of patients (55).

In our proof-of-concept study, two negative samples, according to our described inclusion criteria for negative samples, were marked positive by two dogs; the information was immediately sent to the concerned hospital through the anonymous samples codes, and PCRs that were redone on the patients gave positive results. As said, the dogs are sniffing for specific molecules induced by the SARS-CoV-2 virus presence and dogs have no reason to lie as long as there is no environmental disturbance during their work.

By registering all the events, data, and filming every single trial for each dog, we were able to understand the reasons for some of the rare false negative marking that occurred: a horse passing closely by the working room (Alfort is a veterinary school…) or the presence of a too zealous television team that did not respect the edited procedures for example.

Sniffing behaviour of the dog is actively controlled during investigatory behaviour and rapidly modulated in response to sensory input so that the transport of VOCs to the olfactory epithelium and thus the olfactory proceeding is optimized (57).

Another case of false positive marking in our study looks interesting to analyse. We took also negative samples of axillary sweat on some the people directly involved in our study. One of these samples, made on a young woman, was marked positive by two male dogs without any environmental disturbance, but the dogs showing more excitation for the sample than usually for other positive ones. We investigated the case and found that this woman had been sampled during her fertile period, and therefor we looked at the sexual pheromones related to the period.

Androsterone, androstenol and isovaleric acid contribute to odour profiles, and the human axillary odour seems to have a communicative role analogous to that of the odours from specialized skin glands of animals. Some animal species have apocrine sudoral or aprocrine sebaceous glands, similar to those of the human axilla. These glands have prominent social functions, including a complex system of chemical messages (pheromones) that provoke specific types of behaviour. Published studies (58) show that the presence of androstenone in female axillary sweat make females more sexually attractive in different species including humans.

In Shakespeare’s day, a woman in her fertile period used to hold a peeled apple under her arm until the fruit became saturated with her armpit scent; then she presented this “love apple” to her lover to inhale in order to provoke his sexual excitation. Pheromones are defined as substances produced by on animal which conveys information to other individuals by olfactory means (59). And in such a situation androsterone and molecules like benzoate derivates, excreted in axillary sweat, enhance sexual attraction by men toward women near the end of follicular phase of the menstrual cycle when fertility is at highest (60, 61).

In bitches, during oestrus, these molecules are part of the secretions that are highly attractive to male dogs and enhance sexual excitement (62, 63, 64). This fact explains the fail (false positive) of the two male dogs, and shows that such situations should be included in future male dogs training on human axillary sweat samples.

In our study, like in many others conducted on dog olfactive detection, the performance is defined in accordance with what is called the signal-detection theory (65, 66). Concha (67) describes it as follow:

1. True positive: the dog indicates the target odour by a “sit” response
2. False positive: the dog alerts to a non target position (control)
3. False negative: the dog fails to exhibit the trained alert in the presence of the target odour
4. True negative: the dog does not alert in the absence of the target odour

All the trials of the dogs were filmed, in order to check afterwards more precisely their sniffing behaviour. This will allow us to determinate the duration of each trial before the dog alerts. The encoding of the presence/absence encoding stimulus is very rapid in the dog, and its discrimination seems to require only one sniff (68), therefor, thinking about future practical applications of our study, prolonged sniffing does not seem to be necessary when the target odour is absent (COVID-19 negative samples or people) (69).

Future analysis of the results will also allow us to have a better appreciation of the impact of the results of the impact of the quality of the samples on the olfactive performances of the dogs. We will also compare the marking behaviours of the dogs with the characteristics of the sampled COVID-19+ persons (age, gender, ethnic origin).

We will work on some limits we are aware of, like the eventual presence in the organism of the COVID-19 positive patients of other pathogens, the dogs’ attitude facing other types of respiratory diseases, and the correlations that might exist between the dog alert on positive samples and other biological tests than RT-PCR.

Finally, after this working period, we also can conclude what will look at what is an evidence for good cynotechnicians: dogs already trained to work on a line of samples and explosives/detection dogs can be efficient to detect COVID-19 very quickly (a few days), where search and rescue dogs need first to be readapted to a new methodology (for them) and will take a few weeks to be efficient.

We conclude that there is a high evidence that dogs can detect a person infected by the virus responsible for COVID-19 disease.

## Conclusion

In a context where, in many countries worldwide, diagnostic tests are lacking in order to set up a mass detection of COVID-19 contaminant people, we think it is important to explore the possibility of introducing dog olfactive detection as a rapid, reliable and cheap “tool” to either pre-test willing people or be a fast checking option in certain circumstances. The first step for such an approach was to determinate if the samples we decided to choose (axillary sweat) could allow the dogs to olfactively discriminate between positive and negative people regarding COVID-19. In this proof-of-concept study provide evidence according to which the axillary sweat of SARS-CoV-2 contaminated persons can be detected by trained dogs.

The next step is to carry out a validation study with the same dogs of this proof-of-concept study which will provide the sensibility and specificity of the dog’s diagnostic. If such sensitivity and specificity are high enough, then this new study will provide evidence that national authorities may use trained dogs to detect COVID-19 in settings where equipment and money are lacking to perform standard serology or RT-PCR tests, or as a complementary method in other settings.

This first study is fully responding to the concept of “one health-one medicine”, as involving medical doctors, veterinarians, dog handlers, chemists and … dogs !

## References

1. Pirrone F, Albertini M. Olfactory detection of cancer by trained sniffer dogs: a systematic review of the literature. J. Vet. Behav. 2017, 19: 105–117

2. Hasell J. Testing early, testing late: four countries’ approaches to COVID-19 testing compared. Ourworldindate.org; 2020, may 19. https://ourworldindata.org/covid-testing-us-uk-korea-italy

3. Van DEN BERG T. The role of the legal and illegal trade of live birds and avian products in the spread of avian influenza. Rev. Sci. Tech. Off. Int. Epiz. 2009; 28(1):93–111

4. Farabee MJ. The integumentary system. In: Farabee MJ, editor. The On-Line Biology Book. 9th edn. Estrella Mountain Community College; Avondale, AZ (USA): 2007; pp. 1–3.

5. Wysocki CJ, Pretti G. Facts, fallacies, fears and frustrations with human pheromones. Anat. Rec. A Discov Mol Cell Evol Biol. 2004; 281(1):1201–1211

6. Troccaz M, Starkenmann C, Niclass Y. 3-Methyl-3-sulfanylhexan-1-ol as a Major Descriptor for the Human Axilla-Sweat Odour Profile. Chem Biodivers. 2004, 1:1022–1035

7. Hasegawa Y; Yabuki M, Matsukane M. Identification of New Odoriferous Compounds in Human Axillary Sweat. Chem. Biodivers. 2004;1(12)2042–2050

8. Zeng XN, Leyden JJ, Lawley HJ. Analysis of the characteristic odors from human male axillae. J. Chem. Ecol. 1991; 17: 1469–1492.

9. Zeng XN, Leyden JJ, Speilman AI, Preti G. Analysis of the characteristic human female axillary odors: qualitative comparison to males. J. Chem. Ecol. 1996; 22; 237–257

10. Jenkins SH. Can police dogs identify criminal suspects by small ? Using experiments to test hypothesis about animal behaviour. In: Jenkins SH.editor. How science works: Evaluating evidence in biology and medicine. Oxford Univ. Press, New York, 2004, pp 36–51

11. Bernoier UR, Kline DL, Barnard DL, Schreck CE, Ost RA. Analysis of human skin emanations by gas chromatography/mass spectrometry. Anal. Chem. 2000; Feb 15; 72(4): 747–756

12. Gallagher M, Wysocki CJ, Leyden JJ, Spiezman AL, Sun X, Preti G. Analyses of volatile organic compounds from human skin. Br. J. Dermatol 2008; Spet; 159(4): 780–791

13. Bijland LA, Bomers MK, Smulders YM. Smelling the diagnosis: a review on the use of scent in diagnosing disease. Neth. J. Med. 2013;71(6):300–307

14. Angle C, Waggoner LP, Ferrando A, Haney P, Passler T. Canine Detection of the Volatilome: A Review of Implications for Pathogen and Disease Detection. Front. Vet. Sci. 2016; june 24; 3:47

15. Williams H, Pembroke A. Sniffer dog in the melanoma clinic ? Lancet. 1989; 1:734

16. Willis CM, Britton LE, Wallace J, Guest CM. Volatile organic compounds as biomarkers of bladder cancer: sensitivity and specificity using trained sniffer dogs. Cancer Biomark. 2010; 11; 8:145–153

17. Willis CM, Church SM; Guest CM; Cook WA, Mccarthy N, Branbury AJ, Church MR, Church JC. Olfactory detection of human bladder cancer by dogs: proof of principle study. BMJ. 2004;329(7468):712

18. Sonoda H, Kohnoe S, Yamazato T, Satoh Y, Morizono G, Shikata K, Morita M, Watanabe A, Morita M, Kakeji Y, Inoue F, Maehara Y. Colorectal cancer screening with odour material by canine scent detection. Gut. 2011;60(6):814–819

19. Sarkis R, Khazen J, Issa M, Khazzala A, Hilal G, Grandjean D. Dépistage précoce du cancer colorectal par detection olfactive canine. Proceedings JFHOD; Paris, France, 22-25/08/2018

20. Boedecker E, Friebel G, Walles T. Sniffer dogs as part of a bimodal bionic research. Approach to develop a lung cancer screening. Interact. Cardiovasc. Thorac Surg. 2012;14(5):511–515

21. Buszewski B, Ligor T, Jezierski T. Wenda-PIESIK A, Walczak M, Rudnicka J. Identification of volatile lung cancer marker by gas chromatography-mass spectrometry: comparison with discrimination by canines. Anal. Bioanal. Chem. 2012; 404(1)141–146

22. Ehmann R., Boedeker E., Friedrich U. Sagert J, Dippon J, Friedel G, Walles T. CANINE scent detection in the diagnosis of lung cancer: revisiting a puzzling phenomenon. Eur. Resp. J. 2012; 39(3);669–676

23. Guirao A, Molins L, Ramon I, Sunyer G, Vinolas N, Marrades R, Sanchez D. et al Trained dogs can identify malignant solitary pulmonary nodules in exhaled gas. Lung Cancer. 2019;135:230–233

24. Pickel D, Manuly GP, Walker DB, Evidence for canine olfactory detection of melanoma. Applied an. Behav. Sc. 2004; 89(1):107–116

25. Campbell LF, Farmery L, Creighton GEORGE SM, Farrant P. Canine olfactory detection of malignant melanoma. BMJ Case Rep. 2013; Published online doi:10.1136/bcr-2013-008566

26. Bjartell AS. Dogs sniffing urine: a future diagnostic tool or a way to identify new prostate cancer markers ? Eur Urol. 2011;59(2):202–203

27. Cornu JN, Cancel-TASSIN G., Ondet V, Girarbet C. Cussenot O. Olfactory detection of prostate cancer by dogs sniffing urine: a step forward in early diagnosis. Eur. Urol. 2011;59(2):197–201

28. Taverna G., Tidu L, Grizzli F. Olfactory system of highly trained dogs detects prostate cancer in urine samples. J. Urol. 2015; 193(4);1382–1387

29. Kitiakara T, Redmond S, Unwanatham N, Rattanasiri S, Thakkinstian A, Tangtawee P, Mingphruedhi S, Sobhonslidsuk A, Intaraprasong P, Kositchaiwat C. The detection of hepatocellular carcinoma (HCC) From patients' breath using canine scent detection: A proof-of-concept study. J. Breath. Res. 2017,11(4):046002

30. Wells DL, Lawson SW, Siriwardena AN, Canine response to hypoglycaemia in patients with type I diabetes. J of Altern and Complimentary Med. 2008; 14(10)1235–1241

31. Rooney NJ, Morant S, Guest C, Investigation into the value of trained glycemia alert dogs to clients with type 1 diabetes. PloS One. 2013;8(8):e69921

32. Ronney NJ, Guest C, Swanson L, Morant S; How effective are trained dogs at alerting their owners in blood glycaemic levels ? Variations in performance of glycaemia alert dogs. PloS One. 2019;14(1):p.e0210092

33. Wilson C, Morant S, Kane S, Pesterfield C, Guest C, Rooney NJ. An owner-independent investigation of diabetes alert dog performance. Front. In Vet. Sci 2019;6:91

34. Wallner W., Ellis T. Olfactory detection of Gypsy Moth pheromones and egg masses by domestic canines. Environ. Entomol. 1976; 5:183–186

35. Richards KM, Cotton SJ, Sandemen RM. The use of detector dogs in the diagnosis of nematode infection in sheep feces. J. Vet. Behav. 2008;3:25–31

36. Guest C, Doggett M, Dewhirst S, D’ALESSANDRO U, Kandeh B, Morant SV, Pinder M, Squires C, Logan J, Lindsay SW. Trained dogs identify people with malaria parasites by their odour. Letter in Lancet Infect. Dis. 2019;19(6):578–580

37. Bomers MK, Van AGMAEL MA, Luik H, Van VEEN MC, Vandenbrouck-GRAULS C, Smulders Y. Using a dog’s superior olfactory sensitivity to identify *Clostridium difficile* in stools and patients: proof of principle study. BMJ. 2012;345:e7396

38. Bomers MK, Van AGMAEL MA, Luik H, Van VEEN MC, Vandenbrouck-GRAULS C, Smulders Y. A detection dog to identify patients with Clostridium difficile infection during a hospital outbreak J. of Infection. 2014;69: 456–461

39. Poling A, Weetjens B. Using giant African pouched rats to detect tuberculosis in human sputum samples; Am. J. Trp. Med. Hyg. 2010;83(6):1308–1310

40. Maurer M, Mccullogh M, Willey A, Hirsch W, Dewey D. Detection of *bacteriuria* by canine olfaction. Open form Infect. 2016, 3(2): ofw051

41. Wolfel R, Corman V, Seilmaier M, Zange S, Muller M, Niemeyer D, Jones T, Vollmar P, Rothe C, Hoelscher M, Bleicker T, Brunink S, Schneider J, Ehmann R, Zwirglmaier K, Drosten C, Wendtner C. Virological assessment of hospitalized patients with Covid-19. Nature. 2020; April 1. doi:10.1038/s41586-020-2196-x

42. He X, Lau E, Wu P, Deng X, Wang J, Hao X, Lau Y, Wong J, Guan Y, Tan X, Mo X, Chen Y, Liao B, Chen W, Hu F, Zhang Q, Zhong M, Wu Y, Zhao L, Zhang F, Cowling B, Li F, Leung G. Temporal dynamics in viral shedding and transmissibility of COVID19. Nat. Med. 2020; Apr15; 26, 672–675. doi:10.1038/s41591-020-0869-5

43. Grandjean D, Haak R, Gerritser R, Massey J, Pritchard C, Schuller P, RiviÈRE S. The search and rescue dog; Aniwa ed., Paris (france), 2007, pp296

44. Lay M, Cheng P, Lim W,. Survival of severe acute respiratory syndrome coronavirus. Clin. Infect. Dis. 2005; 41(7):e67–71

45. Shi-YAN R, Wen-BIAO W, Ya-GUANG H, Hao6RAN Z, Zhi-CHAO W, Ye-LIN C, Rong-DING G. Stability and infectivity of coronavirus in inanimate environments. World J. Clin. Cases. 2020; Apr 26; 8(8):1391–1399

46. PRO/AH/EDR>COVID19 update(30): China (Hong Kong), Dog suspected serology pending 2020

47. PRO/AH/EDR>COVID19 update(45): China (Hong Kong), animal, dog; 2^nd^ case PCR positive 2020

48. COVID19 Real PCR validation studies: sequence blast analyses and cross reactivity studies. March 2020. Datas stay by IDEXX laboratories Inc, Westbrook, Maine, USA.

49. Temman S, Barbarino A, Maso D, Dehilil S, Enouf V, Huon C, Eloit M. Absence of SARS-CoV-2 infection in cats and dogs in close contact with a cluster of COVID19 patients in a veterinary campus. bioRxiv; Apr2020, doi: 10-1101/2020.04.07.029090

50. Center for disease control prevention; Coronavirus disease 2019 (COVID19), pets and other animals. https://www.cdc.gov/coronavirus/2019-ncov/animals/pets-other-animals

51. Agence Nationale de Sécurité Environnementale et Sanitaire; Covid19: Domestic animals play no part in transmission of the virus to humans. https://www.anses.fr/en/content/covid-19

52. http://www.randomization.com

53. Sergent E. Calculateurs épidémiologiques Epitool Ausvet ed. Canberra (Australia), 2018, https://epitools.ausvet.com.au/

54. Xiao A, Tong Y, Zhang S. False negative of RT-PCR and prolonged nucleic acid reconversion in COVID-19: rather than recurrence. J. Med. Virol. 2020;10.1002/jnv.25855

55. Yafang L, Lin Y, Lei C, Yiyan S, Zhifang C, Chuhua Y. Stability issues of RT-PCR testing of SARS-CoV-2 for hospitalized patients clinically diagnosed with COVID-19. J. Med. Vorol. 2020;10.1002/jmv.25786

56. Tahamtan A, Ardebili A. RealTime RT-PCR in COVID-19 detection: issues affecting the results. Expert. Rev Mol. Diagn. 2020; 20(5):453–454

57. Wachowiak M; All in a sniff: olfaction as a model for active sensing. Neuron. 2011;71:962–973

58. Labows JN, Mc GINLEY KJ, Kligman AM. Perspectives on axillary odor. J. Soc. Cosmet. Chem. 1982, 34:193–202

59. Albone E. Mammalian semiochemistry: the investigation of chemical signals between mammals; 1984; Winley ed., New York (USA) pp280

60. Williams M, Jacobson A. Effect of copulins on rating of female attractiveness, mate-guarding, and self-perceived sexual desirability. Evol. Psychol. 2016;14(2):1–8

61. Cherry J, Baum M. Sex differences in main olfactory system pathways involved in psychosexual function. Genes, Brain and Behav. 2020;19:e12618

62. Goodwin M, Gooding K, Reginer F. Sex pheromone in the dog. Science. 1979, 203:559–561

63. Bekoff M. Scent marking by free ranging domestic dogs: olfactory and visual components. Biol. Behv 1979; 4:123–139

64. Macdonald DW. The carnivores: Order carnivora. In: Social odours in mammals Vol.2; Brown RE and MacDonald DW Eds; Clarendon Press, Oxford (UK); 1985; pp619–722

65. Bjellanger R, Andersen E, Maclean I. A training program for filter-search mine detection dog. Int. J. Comp Psych. 2002, 15:278–287

66. Macmillann N, Creezman C. Basic detection theory and one-interval design. In: MacMillann, N, Creezman C Ed. Detection theory: a user’s guide 2^nd^ ed. Laurence Erlbaum Associates; 2005:3–25.

67. Concha A, Mills D, Feugier A, Zulch, Guest C, Harris R, Pike T. Using sniffing behaviour to differenciate true negative from false negative responses in trained scent-detection dogs. Chem. Senses; 2014; 39:749–754

68. Mainland J., SOBEL N; The sniff is part of the olfactory perception. Chem. Senses; 2006;31:181–196

69. Wesson D, Verhagen J, Wachowiak M. Why sniff fast ? The relationship between sniff frequency, odor discrimination, and receptor neuron activation in the rat. J. Neurophysiol. 2009;101:1089–1102

70. Kirton A, Winter A, Wirrel E. Seizure response dogs: evaluation of a formal training progress. Epilepsy and Behaviour. 2008;13(3):499–504

